# Molecular identification of vertebrate blood meals in *Culicoides* biting midges reveals host-vector associations relevant to arbovirus transmission in Ireland

**DOI:** 10.64898/2026.09.10.750564

**Authors:** Elsie Isiye, Angela Valcarcel Olmeda, David O’Neill, James O’Shaughnessy, Annetta Zintl, Denise O’Meara

**Author notes:** Correspondence: Elsie Isiye. Joint last authors (Annetta Zintl, Denise O’Meara).

## Abstract

Understanding the vertebrate hosts exploited by *Culicoides* biting midges is fundamental to assessing the transmission risk of livestock arboviruses. Blood meal analysis provides direct evidence of host-vector interactions under natural conditions; however, molecular data on *Culicoides* host feeding are lacking in Ireland. This study aimed to identify the vertebrate blood meal hosts of *Culicoides* species of veterinary importance and establish host-vector associations relevant to arbovirus transmission.

A total of 115 morphologically blood-fed *Culicoides* females collected from 13 sites across Ireland were analysed using a sequential molecular workflow. Species-specific real-time PCR assays were used to screen for vertebrate hosts, with positive detections confirmed by mitochondrial DNA sequencing. Blood-fed *Culicoides* were subsequently identified to species level using DNA barcoding, enabling direct linkage between vector species and their vertebrate hosts. Vertebrate DNA was detected in 50 specimens, with cattle and horses accounting for the majority of identified blood meals. *Culicoides obsoletus* accounted for most cattle (n = 12) and horse (n = 11) blood meals. *Culicoides scoticus* fed on cattle (n = 6), horses (n = 4) and sheep (n = 1), while *C. dewulfi* and *C. chiopterus* fed predominantly on cattle, with occasional horse blood meals. Four cattle blood meals were identified from *C. punctatus*. No cervid-derived blood meals were detected. This study provides the first molecular characterisation of *Culicoides* blood meal hosts in Ireland, confirming host-vector associations between major vector species and livestock hosts of veterinary importance. Although the number of successfully characterised blood meals was modest, reflecting the inherent difficulty of recovering recently blood-fed *Culicoides* from field collections, the findings establish an important baseline for Ireland. The sequential molecular workflow provides a practical framework for future blood meal investigations and will support improved understanding of host-vector interactions relevant to arbovirus surveillance and risk assessment.

## Introduction

*Culicoides* biting midges are recognised vectors of important livestock arboviruses, including bluetongue virus (BTV) and Schmallenberg virus (SBV), both of which have emerged across Europe in recent decades (Carpenter et al. 2009; 2013). Transmission occurs during blood feeding, when female *Culicoides* obtain vertebrate blood meals required for egg maturation (Purse et al. 2015). Consequently, the epidemiological signifcance of these vectors depends largely on their feeding interactions with susceptible vertebrate hosts. Blood meal analysis provides the most direct approach for identifying hosts utilised by *Culicoides* under natural conditions through the analysis of residual vertebrate blood within the abdomen of blood-fed females (Martínez-de La Puente et al. 2015; Kasičová et al. 2021; González et al. 2022). Characterising these host-vector associations is fundamental to understanding pathogen transmission dynamics and assessing arbovirus risk.

Although *Culicoides* species are broadly classified as either mammophilic or ornithophilic, host selection is often influenced more by host availability and accessibility than by strict host preference (Bartsch et al. 2009; Garros et al. 2011; Slama et al. 2015). Many vector species display generalist feeding behaviour, exploiting multiple vertebrate hosts under natural conditions (England et al. 2020; Martínez-de La Puente et al. 2015). Mammophilic species are of greatest veterinary importance, with cattle, sheep, horses, and deer commonly identified as blood meal hosts, whereas ornithophilic species are also widespread and, in some cases, exhibit strong avian host specificity (Černý et al. 2012; Ninio et al. 2011; Lassen et al. 2012; Martínez-De La Puente et al. 2011). To better understand these host-feeding patterns, molecular approaches have increasingly been applied to improve the sensitivity and taxonomic resolution of host identification, enabling more detailed investigations of vector-host interactions across diverse *Culicoides* species (Calvo et al. 2012; Martínez-de La Puente et al. 2015; Ogola et al. 2023; Tomazatos et al. 2020). In particular, mitochondrial markers such as cytochrome b and cytochrome c oxidase subunit 1 (CO1) are commonly used because of their high copy number and suitability for analysing degraded samples (Alcaide et al. 2009; Hadj-Henni et al. 2015; Kocher et al. 1989; Lassen et al. 2012; Reeves et al. 2018). However, the success of host identification can vary considerably across studies, as it is influenced by factors including the timing of specimen collection, the stage of blood meal digestion, sample storage and preservation conditions, and the specificity of molecular markers for resolving vertebrate DNA (Bellekom et al. 2023; England et al. 2020; Kapustová et al. 2025; Kazak et al. 2024).

Despite increasing research on *Culicoides* host use, major knowledge gaps remain in Ireland. Previous outbreaks of *Culicoides*-borne viruses, including SBV (Barrett et al. 2015; Bradshaw et al. 2012; Collins et al. 2019), together with recent detections of BTV in Ireland (Gov.UK, 2025; NDCC Bluetongue Virus Update 1 of 2026), highlight the country’s vulnerability to *Culicoides*-borne arboviruses in the presence of competent vector populations and susceptible vertebrate hosts. National surveillance programmes have successfully documented the distribution of multiple *Culicoides* vector species implicated in SBV and BTV transmission (Collins et al. 2018; Jess et al. 2018; McCarthy et al. 2010; Isiye et al. 2025). However, despite these advances, no study to date has directly identified the vertebrate blood meal hosts of *Culicoides* species in Ireland. Consequently, current assumptions regarding epidemiological risk remain largely inferential, as the presence of a competent vector species alone does not confirm epidemiological relevance unless that species feeds on susceptible or reservoir hosts. Therefore, characterising host-feeding behaviour is essential for assessing the role of Irish *Culicoides* populations in arbovirus transmission and for refining disease risk assessments within livestock production systems. Addressing this knowledge gap formed the basis of the present study to molecularly identify vertebrate blood meal hosts of *Culicoides* spp. in Ireland using a sequential molecular approach involving real-time PCR screening and DNA sequencing, while also identifying vector species to assess host-vector associations. This study provides the first molecular investigation of *Culicoides* host feeding in Ireland while also evaluating the performance and limitations of molecular blood meal methodologies.

## Methods

### Morphological selection

*Culicoides* populations were monitored at 13 study sites across Ireland using standardised entomological sampling with Onderstepoort Veterinary Institute (OVI) light-suction traps between June 2022 and December 2023. Following collection, specimens were preserved in 70% ethanol. In the laboratory, female *Culicoides* were observed under a stereomicroscope and classified according to their abdominal pigmentation. Females exhibiting pigmented abdomens (parous individuals) or containing visible blood meals were retained for further assessment, whereas females with unpigmented abdomens, indicative of nulliparity, were excluded from subsequent analyses. The degree of blood digestion was evaluated visually to develop a pictorial reference ensuring consistency and standardisation across samples selected where only blood-fed females containing comparable stages of digestion were selected for downstream molecular analyses. Nulliparous females were characterised by pale, translucent abdomens with no visible blood content while blood-fed individuals exhibited progressive changes in abdominal distension and pigmentation consistent with blood meal digestion (Fig. 1).

**Fig. 1.**
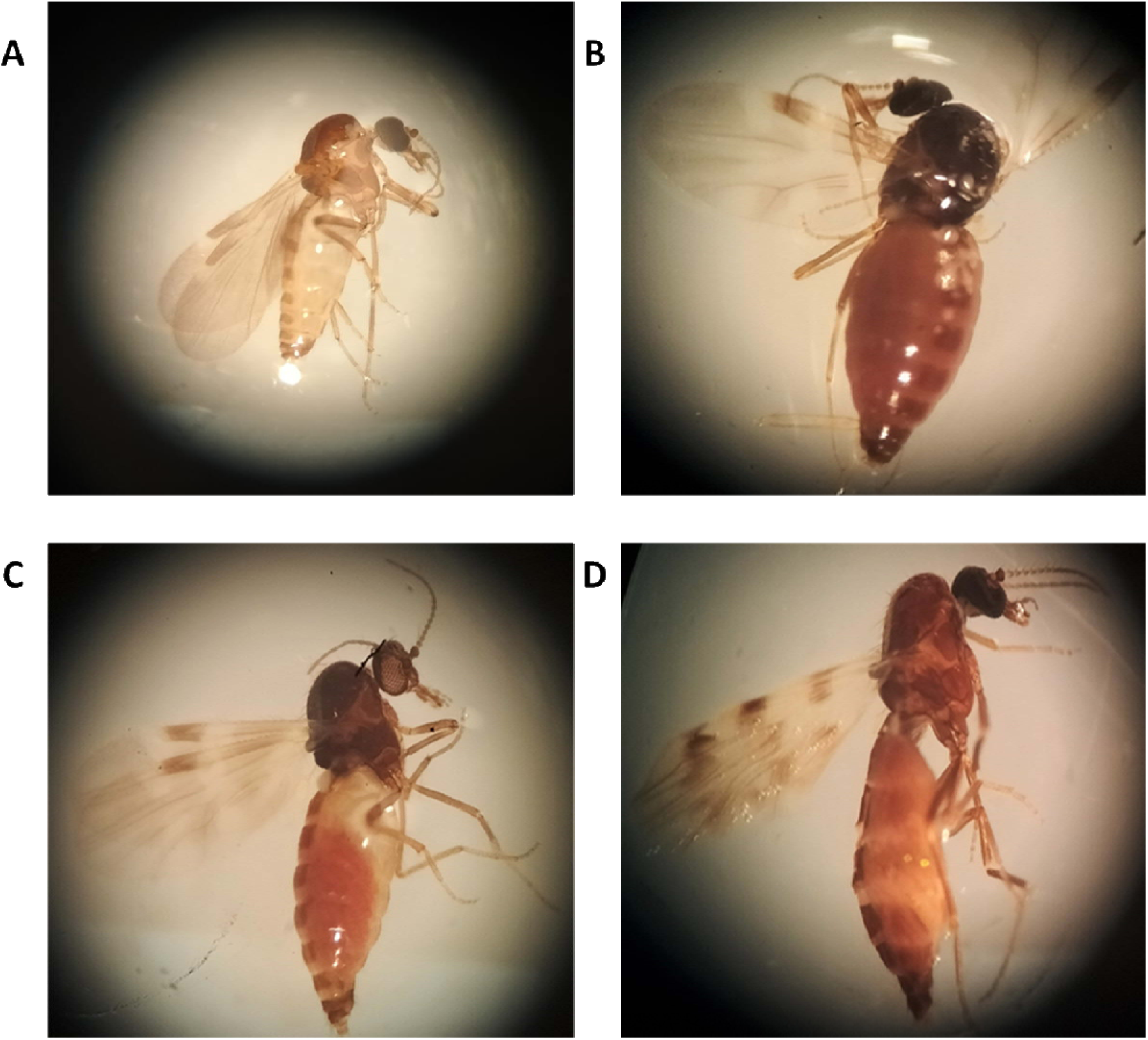
Classification of blood meal digestion stages in female *Culicoides*. Representative examples show (A) unfed individual (B) fully engorged individual with distended abdomen and intense reddish-brown pigmentation, (C) early digestion stage with partial distension and reduced pigmentation, and (D) advanced digestion stage with faint brown discolouration

Three blood-feeding stages were distinguished morphologically: fully engorged females showed markedly swollen abdomens with reddish to dark brown pigmentation indicative of recent feeding; early-stage digesting individuals displayed moderate distension with reduced coloration; and late-stage digesting specimens had contracted abdomens with faint brownish pigmentation, reflecting advanced blood degradation (Fig. 1).

### Real-time PCR screening of vertebrate hosts

Each specimen was crushed using a pipette tip in a sterile 1.5 mL microcentrifuge tube and DNA was extracted using the NucleoSpin DNA Extraction Kit (Macherey-Nagel) according to the manufacturer’s instructions. Final DNA elution was performed in 30 µL of elution buffer and DNA concentration and purity were assessed using a NanoDrop ND-8000 spectrophotometer. Species-specific real-time PCR primer sets targeting conserved mitochondrial regions were obtained from published studies and selected to enable the rapid detection of the veterinary important vertebrates in Ireland (Table 1). These include assays for cattle (*Bos taurus*), sheep (*Ovis aries*) (Dooley et al. 2004), horses (*Equus caballus*) (Meira et al. 2016), and deer species - red (*Cervus elaphus*), fallow (*Dama dama*), and sika (*Cervus nippon*) (Powell et al. 2015). This screening approach was used to prioritise samples for confirmatory DNA barcoding. Real-time PCR runs were performed for each extracted DNA from blood-fed female *Culicoides* to screen for the presence of vertebrate host DNA. Each reaction was performed in a final volume of 10 µL, consisting of 5 µL of 2× FastStart Universal SYBR Green Master Mix (ROX) (Sigma-Aldrich), 0.4 µL of combined forward and reverse primers (final concentration 200 nM each) (Table 1), 3.6 µL of molecular-grade water, and 1 µL of extracted DNA as the template. No template control (NTC) reactions included 1 µL of molecular-grade water in place of the DNA template. Positive control DNA extracts from cattle, sheep, horse, fallow deer, sika deer, and red deer were used to validate the primer performance and assay sensitivity. DNA samples were initially screened in single reactions, and those showing amplification were subsequently analysed in triplicate to confirm reproducibility and to provide an additional quality control measure.

**Table 1.**
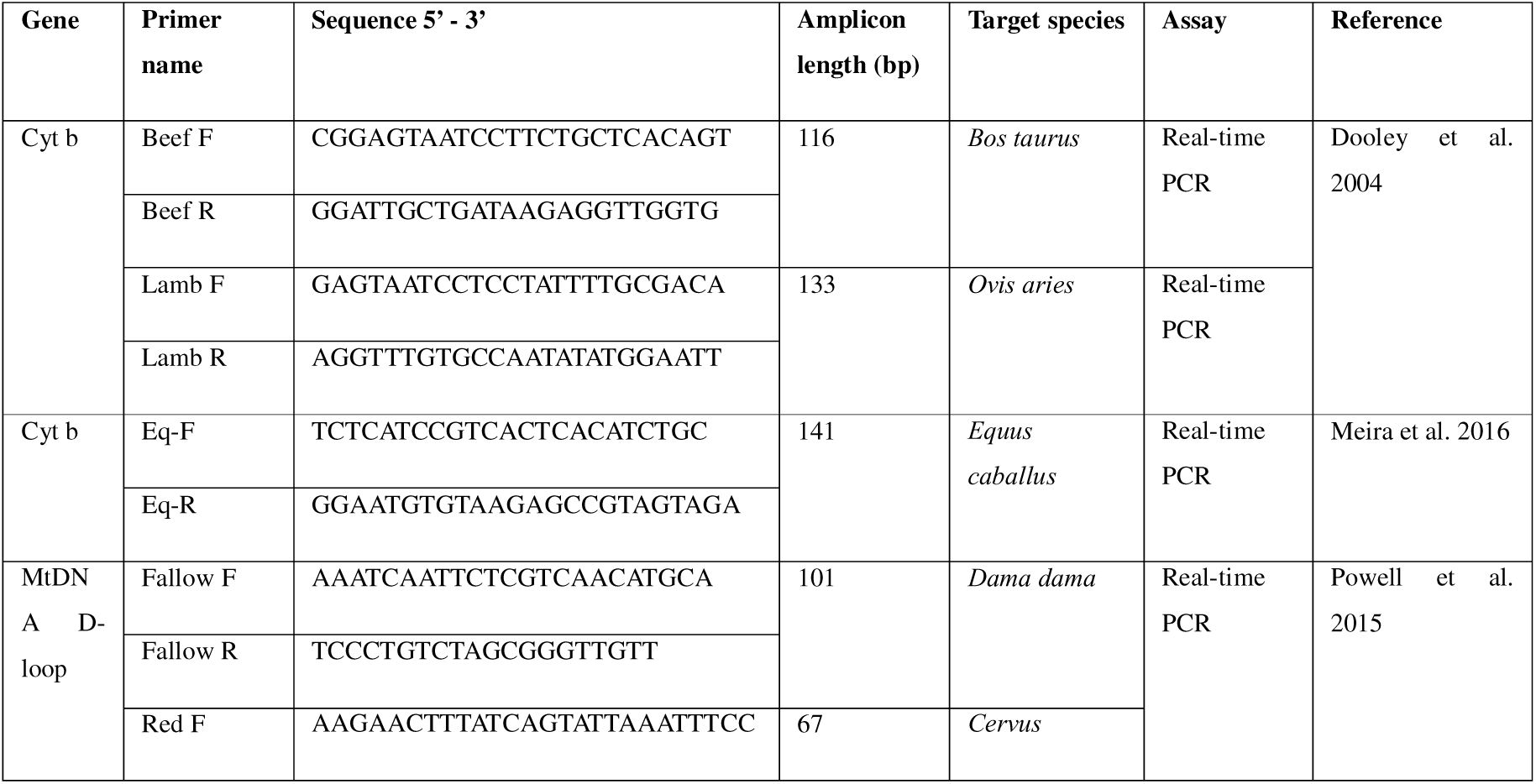
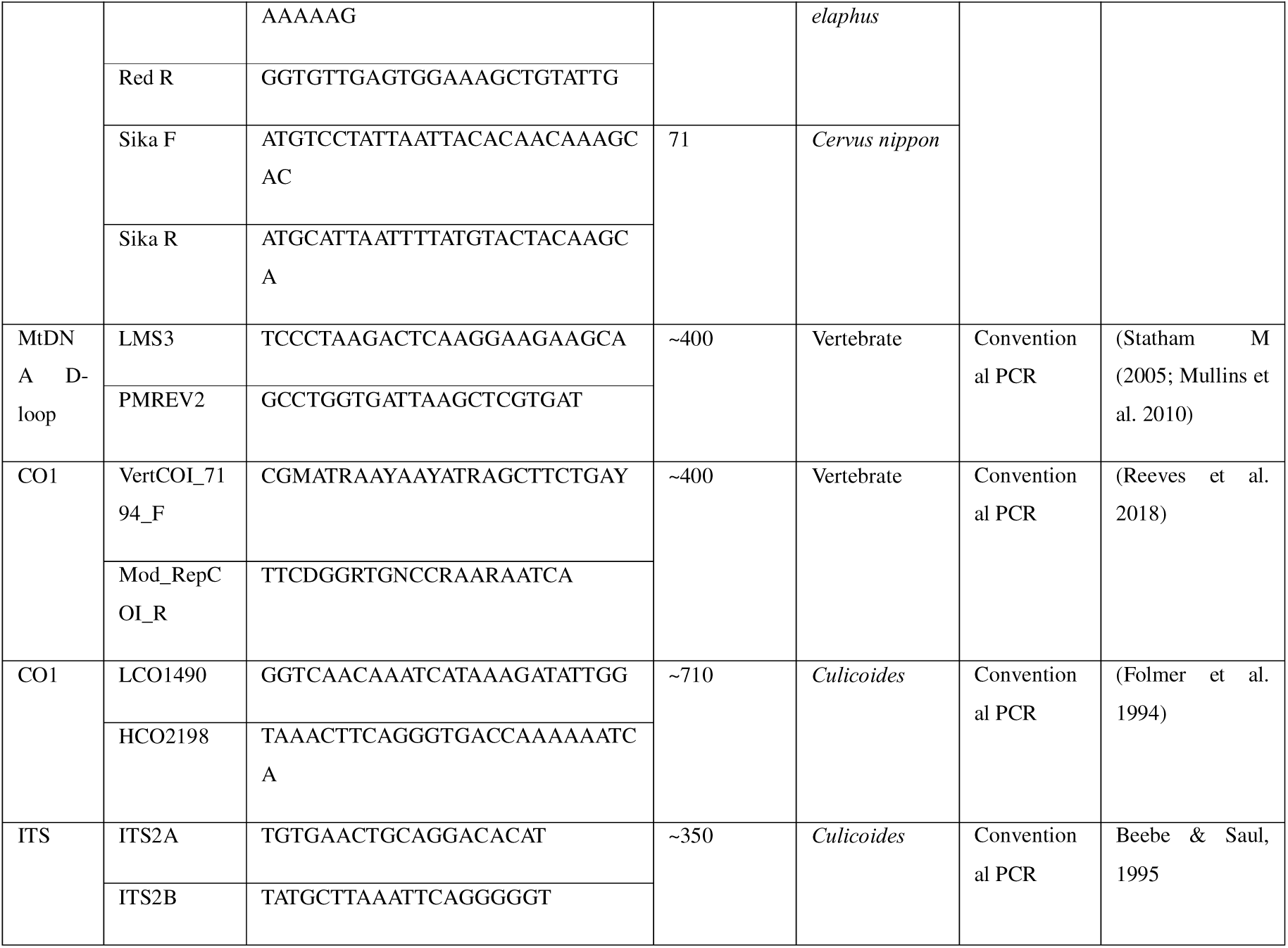
Primers used to identify vertebrate hosts and *Culicoides* vector species.

### DNA sequencing confirmation of vertebrate hosts

DNA sequencing of vertebrate mitochondrial markers was subsequently performed to confirm host identity. Conventional PCR amplification targeted the mitochondrial (D-loop) region using the primer pair LMS3/PMREV2 (Table 1), which has been shown to amplify approximately 400 bp of the mitochondrial d-loop region of a broad range of vertebrate taxa (Statham, 2005; Mullins et al. 2010). In silico BLAST analysis was performed on these primers to assess potential amplification across vertebrate taxa. PCR reactions were performed in a final volume of 10 µL, containing 5 µL of DreamTaq Green Master Mix (Promega), 2 µM of each primer, 1 µL of template DNA, and molecular-grade water. Thermal cycling conditions consisted of an initial denaturation at 95 °C for 2 min, followed by 50 cycles of denaturation at 95 °C for 30 s, annealing at 57 °C for 30 s, and extension at 72 °C for 30 s, with a final extension at 72 °C for 10 min and a 4 °C hold, as described by Statham (2005). In addition, an alternative COI primer pair (VertCOI_7194_F and Mod_RepCOI_R), also amplifying an approximately 400 bp of vertebrate mitochondrial DNA (Reeves et al. 2018; Ogola et al. 2024) was applied. PCR reactions were carried out in a final volume of 20 µL, containing 10 µL of DreamTaq Green Master Mix, 0.75 µL of each primer (10 µM), 7.5 µL of sterile molecular-grade water, and 1 µL of the extracted DNA. The thermocycling conditions included an initial denaturation at 94 °C for 3 min, followed by 40 cycles of 94 °C for 40 s, 48.5 °C for 30 s, and 72 °C for 60 s, with a final extension at 72 °C for 7 min and a 4 °C hold. All PCR products were visualised by gel electrophoresis on 1.6% agarose gels and positive amplicons were purified using a microCLEAN purification kit (Microzone Ltd.) following the manufacturer’s instructions. Purified PCR products were sequenced in both forward and reverse directions using an Applied Biosystems 3500 Genetic Analyser using the BigDye Terminator Cycle Sequencing Kit v3.1 (Applied Biosystems). The resulting consensus sequences were then analysed using the nucleotide Basic Local Alignment Search Tool (BLASTn) (Altschul et al. 1990) on GenBank database to confirm species identity.

### *Culicoides* identification

*Culicoides* specimens confirmed to contain a blood meal by at least one of the previously outlined methods were identified to species level. Molecular identification of *Culicoides* specimens was carried out through PCR amplification and sequencing of the mitochondrial cytochrome c oxidase subunit I (COI) gene using the universal invertebrate primers LCO1490/HCO2198 (Folmer et al. 1994) and the internal transcribed spacer 2 (ITS2) region using the ITS2A/ITS2B primers (Beebe & Saul, 1995) (Table 1). These methods enabling species-level identification were as described by Isiye et al. (2025).

### Host-vector association

Host-vector associations identified through molecular analyses were processed and visualised in R version 4.4.1 (R Core Team, 2025). Graphical visualisation was conducted using *ggplot2* (Wickham et al. 2023) and alluvial plots illustrating categorical associations between *Culicoides* species and vertebrate hosts were generated using the *ggalluvial* package (Brunson, 2023). The diagrams depict the proportional contribution of individual *Culicoides* species to different vertebrate host groups, allowing the patterns of host preference, host breadth, and potential bridge-vector roles to be readily observed.

## Results

### Blood-fed specimen retrieval

A total of 115 blood-fed midges, were selected from across the 13 sites and the land use characteristics of each site was recorded in Table 2.

**Table 2.**
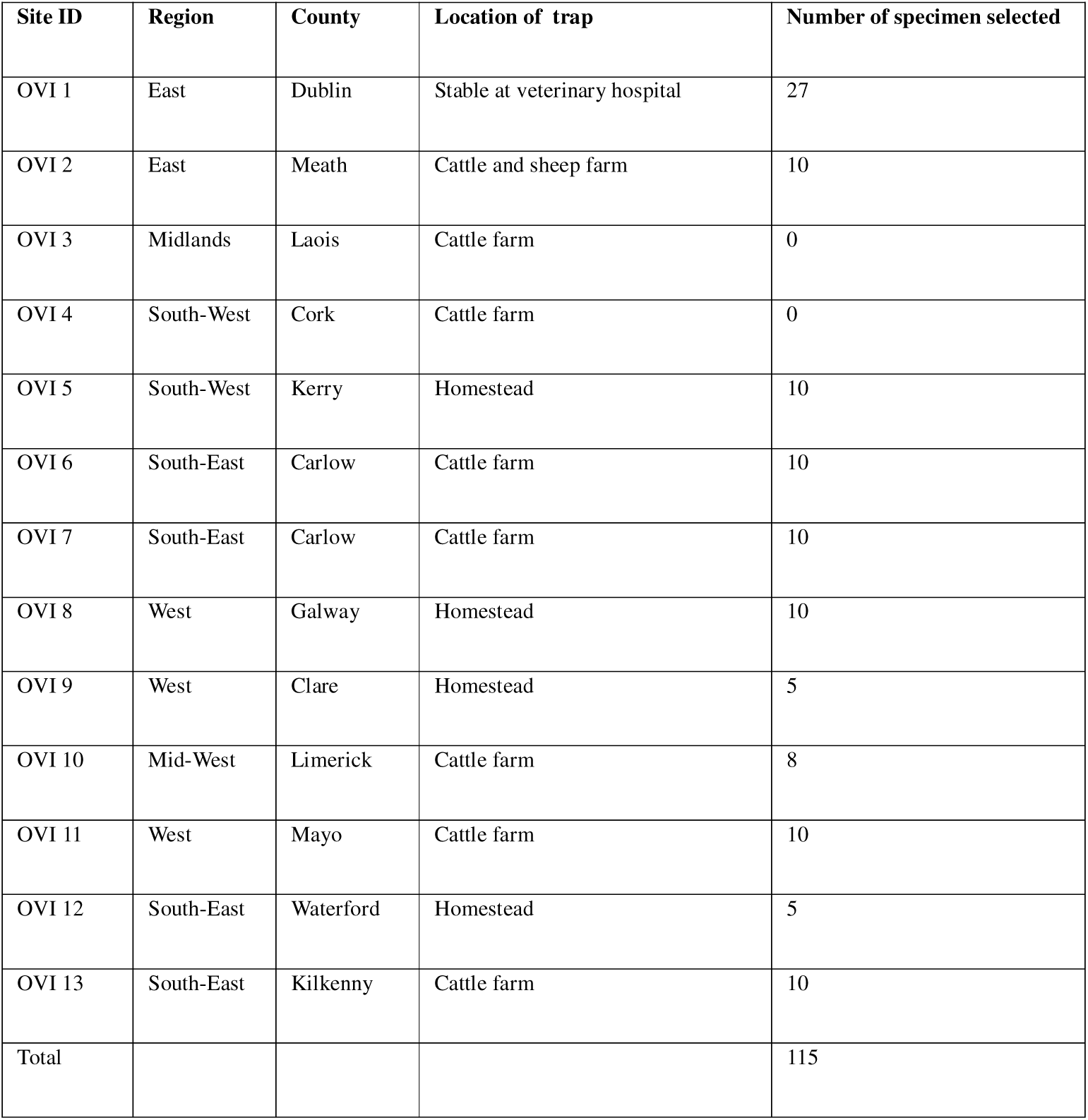
Thirteen trapping locations across Ireland with associated land-use context and the number of morphologically blood-fed female midges selected for downstream molecular analyses. Blood-fed specimens were defined as those with visible reddish or brown abdominal pigmentation consistent with ingested vertebrate blood.

### DNA extraction and PCR amplification

DNA yields obtained from homogenised blood-fed *Culicoides* individuals ranged from 4 to 25 ng/µl. Successful amplification was characterised by the consistent amplification across triplicate reactions (Fig. 2) supporting the reproducibility of target detection and lack of amplification of the NTCs minimised the likelihood of false-positives.

**Fig. 2.**
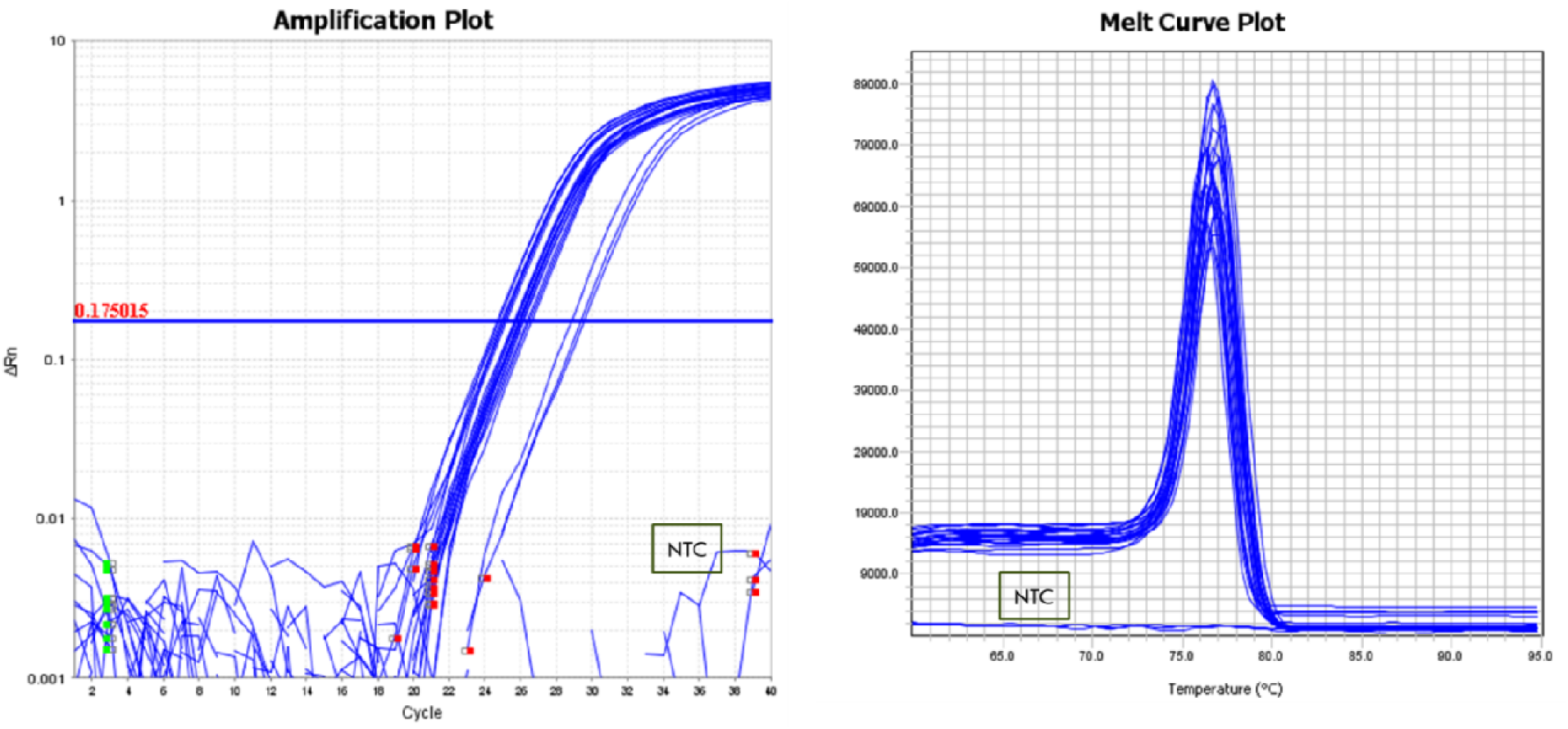
Triplicate real-time PCR confirmation of equine-derived blood meals in *Culicoides* specimens. Consistent amplification across replicates confirmed reproducibility.

A total of 50 specimens tested positive for vertebrate host DNA using species-specific assays. These included 31 bovine-positive samples, 18 equine-positive samples, and one ovine-positive sample. No cervid DNA was detected in any analysed specimens (Supplementary Material S1)

Using the LMS3/PMREV2 primers, vertebrate DNA amplification was successful, producing PCR products of approximately 400 bp. Sequencing analysis identified one specimen as *Ovis aries*, while 28 specimens were confirmed as *Bos taurus* through conventional PCR and subsequent sequencing. Consistent with in silico BLAST predictions (Supplementary Material S2), equine DNA was not amplified with the LMS3/PMREV2 primers, as equids were not represented within the predicted primer target range. The VertCOI_7194_F/Mod_RepCOI_R primer set produced amplicons of approximately 400 bp, and sequencing successfully confirmed five samples as *Equus caballus*. However, sequencing of two additional amplicons identified *Culicoides dewulfi* by BLAST analysis, indicating off-target amplification of vector DNA. Nevertheless, the primers provided confirmatory evidence for the presence of equine blood meals in a subset of *Culicoides* specimens. BLAST analysis results for all successfully amplified and sequenced vertebrate DNA obtained using the both vertebrate primer sets are provided in the Supplementary Material S3. Overall, of the 50 blood meals indicated by real-time PCR, DNA sequencing provided confirmatory sequence data for 34 samples (68.0%). There was 100% concordance (n=34) between DNA sequencing results and the original real-time PCR screening results.

### *Culicoides* species identification

All 50 blood-fed *Culicoides* specimens were successfully identified to species level using a combination of CO1 and ITS2 sequencing markers. Of these, 36 specimens were identified using the CO1 marker, while the remaining 14 were resolved using ITS2 sequencing. Species composition was dominated by members of the subgenus *Avaritia*. *Culicoides obsoletus* was the most frequently detected species (n = 23; 46%), followed by *C. scoticus* (n = 11; 22%), *C. dewulfi* (n = 6; 12%), *C. chiopterus* (n = 6; 12%), and *C. punctatus* (n = 4; 8%). BLAST analysis results for all successfully sequenced *Culicoides* specimens using the CO1 and ITS2 primer sets are provided in Supplementary Material S4.

### Host-vector association

The vertebrate species identified by real-time PCR as well as conventional PCR and sequencing are presented in Table 3.

**Table 3.** Vertebrate host species identified from blood-fed *Culicoides* and associated vector species across the study sites. Summary of host identification by real-time PCR and DNA sequencing linked to molecularly identified *Culicoides* species. Dashes indicate samples for which sequencing confirmation was unsuccessful

| Sample site | *Culicoides* species | Vertebrate host (real-time PCR) | Vertebrate host (DNA sequencing) |
| --- | --- | --- | --- |
| OVI 1 | *C. obsoletus* | *Equus caballus* | - |
|  | *C. obsoletus* | *Equus caballus* | - |
|  | *C. chiopterus* | *Equus caballus* | - |
|  | *C. obsoletus* | *Equus caballus* | - |
|  | *C. obsoletus* | *Equus caballus* | *Equus caballus* |
|  | *C. scoticus* | *Equus caballus* | - |
|  | *C. obsoletus* | *Equus caballus* | *Equus caballus* |
|  | *C. scoticus* | *Equus caballus* | - |
|  | *C. obsoletus* | *Equus caballus* | - |
|  | *C. obsoletus* | *Bos taurus* | - |
|  | *C. dewulfi* | *Equus caballus* | - |
|  | *C. dewulfi* | *Equus caballus* | - |
|  | *C. obsoletus* | *Equus caballus* | *Equus caballus* |
|  | *C. obsoletus* | *Equus caballus* | - |
|  | *C. obsoletus* | *Equus caballus* | - |
|  | *C. scoticus* | *Equus caballus* | - |
|  | *C. obsoletus* | *Equus caballus* | *Equus caballus* |
|  | *C. obsoletus* | *Equus caballus* | *Equus caballus* |
|  | *C. scoticus* | *Equus caballus* | - |
| OVI 2 | *C. scoticus* | *Bos taurus* | *Bos taurus* |
|  | *C. chiopterus* | *Bos taurus* | *Bos taurus* |
|  | *C. scoticus* | *Bos taurus* | *Bos taurus* |
|  | *C. chiopterus* | *Bos taurus* | *Bos taurus* |
|  | *C. scoticus* | *Bos taurus* | *Bos taurus* |
|  | *C. punctatus* | *Bos taurus* | *Bos taurus* |
| OVI 5 | *C. scoticus* | *Ovis aries* | *Ovis aries* |
|  | *C. pulicaris* | *Bos taurus* | *Bos taurus* |
|  | *C. punctatus* | *Bos taurus* | *Bos taurus* |
|  | *C. obsoletus* | *Bos taurus* | *Bos taurus* |
| OVI 6 | *C. dewulfi* | *Bos taurus* | *Bos taurus* |
|  | *C. dewulfi* | *Bos taurus* | *Bos taurus* |
| OVI 7 | *C. obsoletus* | *Bos taurus* | *Bos taurus* |
|  | *C. obsoletus* | *Bos taurus* | *Bos taurus* |
|  | *C. dewulfi* | *Bos taurus* | *Bos taurus* |
| OVI 8 | *C. obsoletus* | *Bos taurus* | *Bos taurus* |
|  | *C. scoticus* | *Bos taurus* | *Bos taurus* |
|  | *C. punctatus* | *Bos taurus* | - |
|  | *C. punctatus* | *Bos taurus* | *Bos taurus* |
| OVI 9 | *C. obsoletus* | *Bos taurus* | *Bos taurus* |
| OVI 10 | *C. dewulfi* | *Bos taurus* | *Bos taurus* |
|  | *C. chiopterus* | *Bos taurus* | *Bos taurus* |
| OVI 11 | *C. chiopterus* | *Bos taurus* | - |
|  | *C. chiopterus* | *Bos taurus* | *Bos taurus* |
|  | *C. scoticus* | *Bos taurus* | *Bos taurus* |
|  | *C. obsoletus* | *Bos taurus* | *Bos taurus* |
|  | *C. scoticus* | *Bos taurus* | *Bos taurus* |
| OVI 12 | *C. obsoletus* | *Bos taurus* | *Bos taurus* |
|  | *C. obsoletus* | *Bos taurus* | *Bos taurus* |
| OVI 13 | *C. obsoletus* | *Bos taurus* | *Bos taurus* |
|  | *C. obsoletus* | *Bos taurus* | - |

Analysis of host-vector feeding interactions revealed that *C. obsoletus* accounted for the highest number of identified blood meal sources. This species was associated with 12 bovine (*Bos taurus*) and 11 equine (*Equus caballus*) blood meals, respectively. *C. scoticus* exhibited a broader host-feeding range, with blood meals derived from all three detected host species, including cattle (n = 6), sheep (*Ovis aries*; n = 1), and horses (n = 4). Additional feeding associations were observed for *C. dewulfi*, which fed predominantly on cattle (n = 4) and horses (n = 2). *C. chiopterus* was associated with five bovine and one equine blood meal, while four bovine blood meals were identified from *C. punctatus* specimens (Fig. 3).

**Fig. 3.**
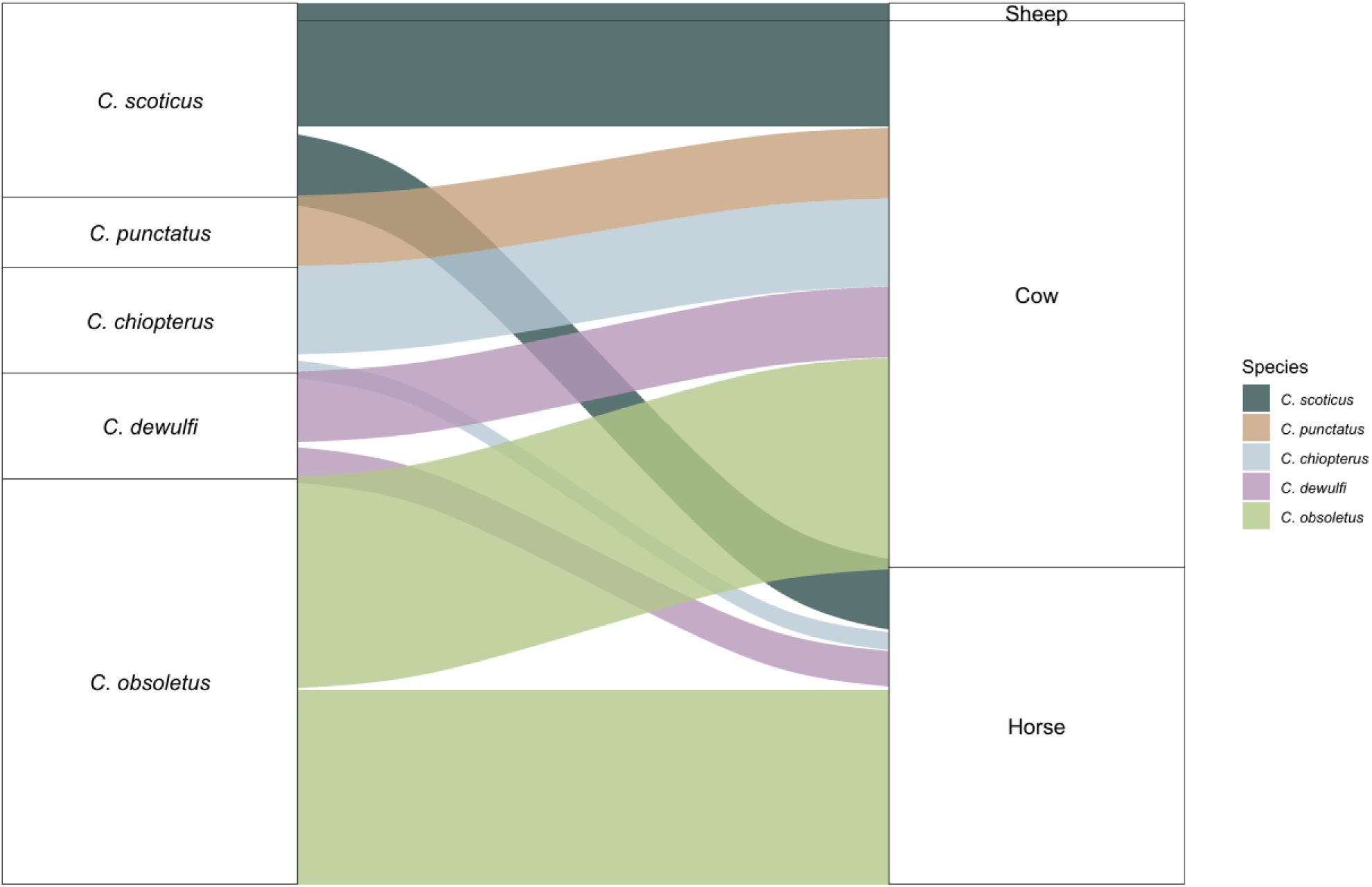
Alluvial diagram illustrating host-vector feeding relationships among blood-fed *Culicoides* species analysed in this study. Flow widths represent the number of confirmed blood meals linking *Culicoides* species to vertebrate hosts (cattle, horses, and sheep), highlighting the dominant feeding associations and host overlap

## Discussion

This study provides the first molecular investigation of *Culicoides* blood meal sources conducted in Ireland and offers new insight into host-vector associations relevant to the transmission of *Culicoides*-borne arboviruses. Across the sampled sites, blood meals were predominantly derived from livestock hosts, with cattle representing the dominant host species identified. Horse and sheep blood meals were also detected, further demonstrating that feeding on domestic livestock species of recognised epidemiological importance. Given that haematophagy is the principal mechanism through which these arboviruses are acquired and transmitted, characterising host-feeding behaviour is essential for understanding transmission dynamics (Carpenter et al. 2009; Martínez-de La Puente et al. 2017; Tarlinton et al. 2012; Wernike et al. 2024). These findings are particularly relevant in the context of SBV and BTV incursions in Ireland and the ongoing risk posed by wind-borne dispersal of infected *Culicoides* into the region (Casey-Bryars et al. 2025; McGrath et al. 2018; Mignotte et al. 2021).

The predominance of members of the Obsoletus complex among blood-fed specimens reinforces the recognised epidemiological importance of these species in Europe. *Culicoides obsoletus* was the most abundant species identified, consistent with previous reports describing it as the dominant *Culicoides* species in Ireland and across much of the Palaearctic region (Isiye et al. 2025; Mignotte et al. 2020). This species showed strong feeding associations with cattle and horses, supporting previous evidence that *C. obsoletus* has substantial ecological plasticity (Martínez-de La Puente et al. 2015; England et al. 2020b). *Culicoides scoticus* displayed the broadest host range among the species analysed, feeding on cattle, sheep, and horses, further supporting its classification as a generalist feeder capable of facilitating cross-species pathogen transmission. Similar feeding behaviour has been reported in previous European studies, which identified both *C. obsoletus* and *C. scoticus* as abundant and epidemiologically important vectors associated with livestock hosts (England et al. 2020; Martínez-de La Puente et al. 2015; Ninio et al. 2011, Elbers & Meiswinkel 2015). The higher detection of blood fed *C. obsoletus* and *C. scoticus* further confirms their epidemiological importance as key vectors of BTV and SBV in Ireland. Other species identified in this study, including *C. dewulfi*, *C. chiopterus*, and *C. punctatus*, fed predominantly on cattle, further highlighting the central role of cattle in the ecology of *Culicoides*-borne viruses in Ireland. The predominance of cattle-derived blood meals was unsurprising given the location of most traps within or adjacent to cattle farming environments; however, it also reflects the recognised attractiveness of cattle to *Culicoides* species and their importance as amplifying hosts for BTV and SBV. Marked variation was observed among trapping sites in the number of blood-fed females recovered. OVI site 1, located within an enclosed stable housing equids on a continuous basis, yielded the highest number of blood fed *Culicoides*. The enclosed indoor environment may have reduced post-feeding dispersal and increased the likelihood of capturing recently engorged females, supporting previous observations that indoor livestock environments can facilitate *Culicoides* feeding activity and resting behaviour (Ninio et al. 2011). Evidence of host-seeking dispersal was also observed at OVI site 2, where *C. chiopterus* specimens containing cattle-derived blood meals were collected despite only sheep being present at the trapping site during sampling. This suggests that individuals may have travelled from neighbouring cattle holdings to acquire blood meals prior to capture, consistent with previous studies demonstrating that *Culicoides* are capable of dispersing several hundred metres between feeding and resting sites (England et al. 2020; González et al. 2022). These observations further highlight the mobility and ecological adaptability of livestock-associated *Culicoides* species within fragmented agricultural landscapes

The methodological approach used in this study represents a significant contribution to *Culicoides* blood meal analysis in Ireland. Initial morphological screening based on abdominal pigmentation and distension provided an efficient means of identifying potentially blood-fed individuals prior to molecular analysis and allowed specimens to be broadly categorised according to digestion stage. This was particularly important given the small blood meal volume ingested by *Culicoides* and the rapid degradation of host DNA during digestion (Chagas et al. 2024; González et al. 2022). The species-specific real-time PCR assays proved highly effective as a primary screening tool, demonstrating high sensitivity and reproducibility for the detection of cattle, sheep, and horse DNA from partially digested blood meals. The higher success of qPCR likely reflects its ability to amplify shorter DNA fragments, making it more suitable for detecting degraded or low-abundance host DNA compared with longer amplicon-based approaches (Keven et al. 2020; Lucero et al. 2014). Previous studies have also demonstrated the effectiveness of real-time PCR in targeted dietary applications where specific host taxa are predefined or cases of degraded templates, such as environmentally-derived DNA (O’Meara et al. 2014) and its application in detecting low-abundance DNA in pooled samples (Isiye et al. 2026). By contrast, high-throughput metagenomic sequencing approaches may be more appropriate where no predefined target species are available (Curran et al. 2022). In this study, real-time PCR therefore represented an appropriate approach given the focus on defined vertebrate hosts and identified host DNA in a greater proportion of samples while sequencing provided confirmation of vertebrate identity. The study also demonstrated important limitations associated with universal primer systems, including occasional amplification of *Culicoides* DNA rather than vertebrate DNA. These findings emphasise the importance of careful primer selection and validation when investigating blood meals from highly degraded samples and demonstrate the trade-offs between sensitivity, specificity, and taxonomic breadth in blood meal analysis methodologies.

Several limitations should be considered when interpreting the findings of this study. First, the number of blood-fed females recovered was relatively low (n = 115 across 13 sites), reflecting a well-recognised limitation of light trap surveillance, which preferentially captures host-seeking rather than recently engorged individuals (Martínez-de La Puente et al. 2015). Digestion stage is a critical factor influencing molecular identification success because host DNA becomes increasingly fragmented as digestion progresses (Reeves et al 2018; Gonzalez et al. 2022). In addition, host availability and trap placement likely influenced the observed feeding patterns. Most traps were positioned within livestock-associated environments, including cattle sheds, milking parlours, and stables, which may have biased collections towards livestock-derived blood meals. The absence of deer-derived blood meals despite the inclusion of cervid-specific assays may therefore reflect limited deer activity near trapping locations, lower local deer abundance, or reduced trapping efficiency within wildlife-associated habitats rather than a true absence of *Culicoides* feeding on cervids. Furthermore, although sample preservation followed standard surveillance procedures, delays between collection, sorting, and DNA extraction may also have contributed to host DNA degradation. Despite these limitations, the inclusive analysis of fully engorged, partially and extensively digested specimens provided a realistic representation of the challenges associated with field-derived *Culicoides* blood meal analysis.

Despite these limitations, this study establishes the first molecular framework for investigating *Culicoides* blood-feeding behaviour in Ireland and provides important baseline data relevant to arbovirus transmission risk. The predominance of cattle-associated blood meals and the dominance of *C. obsoletus* and *C. scoticus* among blood-fed specimens reinforce the epidemiological importance of these species in the transmission of livestock-associated arboviruses. These findings are particularly significant given the recent detection of BTV in Ireland, the continued circulation of BTV in neighbouring regions, and the previous emergence of SBV within Irish livestock populations (Collins et al. 2019; Department for Environment, Food & Rural Affairs and Animal and Plant Health Agency, 2025; NDCC Bluetongue Virus Update 1 of 2026). The relevance of molecular blood-meal identification extends beyond Ireland, particularly in the context of the circulation of these *Culicoides*-borne viruses across Europe. The recent detection of a novel European Shamonda-like Orthobunyavirus in cattle further highlights the need to understand vector-host interactions and potential transmission pathways associated with emerging viruses (Wernike et al., 2026). The molecular blood-meal identification approach presented here therefore provides a transferable tool that could support surveillance and risk assessment of emerging *Culicoides*-borne pathogens across Europe by characterising the host-feeding patterns that underpin vector-host contact.

## Conclusion

This study demonstrates that the integrated use of morphological sorting, real-time PCR screening and DNA sequencing provides a robust framework for characterising host-vector associations in *Culicoides* midges. This approach enabled reliable identification of vertebrate blood meal sources in blood-fed female Culicoides, providing direct evidence of host utilisation patterns. The findings offer a useful methodological framework for future surveillance and the observed associations between vectors and livestock highlight their continued importance within Irish agricultural ecosystems and support their inclusion in ongoing vector monitoring programmes.

Future work should build on this by increasing sample sizes, extending seasonal and spatial coverage, and incorporating trapping strategies that better capture wildlife-derived blood meals. The integration of broader host detection approaches, such as next-generation sequencing, alongside targeted assays will further refine understanding of host range and feeding behaviour. These advances are critical for resolving the complexity of host-vector-pathogen networks and improving risk assessment of *Culicoides*-borne viruses in Ireland and comparable temperate regions.

## Acknowledgments

This research was conducted as part of the Network of Insect Vectors (NetVec) Ireland project (https://www.ucd.ie/netvecireland/), funded by the Department of Agriculture, Food, and the Marine. The authors sincerely thank the NetVec Ireland team and the midge survey volunteers across Ireland for their efforts in specimen collection. The invaluable support and cooperation of these collaborators were instrumental in the successful completion of this study.

## Data availability statements

All data relating to *Culicoides* blood meal analysis that support the findings of this study are included in this article and its Supplementary Material files.

## Author contributions

EI conducted the laboratory work, performed data analysis, and prepared the original and final manuscript. AVO was responsible for specimen collection and sorting. DON provided technical expertise in molecular methods used and contributed to project supervision. JOS contributed midge specimens. AZ provided overall project supervision and led funding acquisition, project conceptualisation, and design. DOM also provided overall project supervision, contributed to project conceptualisation and design, and contributed midge specimens. All authors reviewed and approved the final version of the manuscript.

Supplementary Material S1: Summary of 115 *Culicoides* specimens selected for blood meal analysis, including sample ID, collection location, date of collection, morphological grouping, and specimens testing positive for vertebrate DNA by real-time PCR.

Supplementary Material S2: Potential vertebrates amplified by the primers LMS3/PMREV 2 from in-silico primer blast analysis.

Supplementary Material S3: BLAST analysis results for specimens with detected vertebrate DNA, including sequence length, percentage identity to reference sequences, and the corresponding GenBank accession numbers used for host identification.

Supplementary Material S4: BLAST analysis results for *Culicoides* specimens, including their molecular identification, gene region used for species determination, sequence length, percentage identity to reference sequences, and corresponding GenBank accession numbers used for insect identification.

## References

1. Alcaide, M., Rico, C., Ruiz, S., Soriguer, R., Muñoz, J., & Figuerola, J. (2009). Disentangling vector-borne transmission networks: A universal DNA barcoding method to identify vertebrate hosts from arthropod bloodmeals. PLoS ONE, 4(9), e7092. 10.1371/journal.pone.0007092

2. Altschul, S. F., Gish, W., Miller, W., Myers, E. W., & Lipman, D. J. (1990). Basic local alignment search tool. Journal of Molecular Biology, 215(3), 403–410. 10.1016/S0022-2836(05)80360-2

3. Augot, D., Hadj-Henni, L., Strutz, S. E., Slama, D., Millot, C., Depaquit, J., & Millot, J.-M. (2017). Association between host species choice and morphological characters of main sensory structures of *Culicoides* in the Palaearctic region. PeerJ, 5, e3478. 10.7717/peerj.3478

4. Bartsch, S., Bauer, B., Wiemann, A., Clausen, P.-H., & Steuber, S. (2009). Feeding patterns of biting midges of the *Culicoides obsoletus* and *Culicoides pulicaris* groups on selected farms in Brandenburg, Germany. Parasitology Research, 105(2), 373–380. 10.1007/s00436-009-1408-y

5. Beebe, N. W., & Saul, A. (1995). Discrimination of all members of the *Anopheles punctulatus* complex by polymerase chain reaction-restriction fragment length polymorphism analysis. The American Journal of Tropical Medicine and Hygiene, 53(5), 478–481.

6. Bellekom, B., Bailey, A., England, M., Langlands, Z., Lewis, O. T., & Hackett, T. D. (2023). Effects of storage conditions and digestion time on DNA amplification of biting midge (*Culicoides*) blood meals. Parasites & Vectors, 16(1), 1–11. 10.1186/s13071-022-05607-x

7. Brunson, J. C. (2023). *ggalluvial: Alluvial plots in ggplot2* (Version 0.12.5) [R package]. https://CRAN.R-project.org/package=ggalluvial

8. Calvo, J. H., Berzal, B., Calvete, C., Miranda, M. A., Estrada, R., & Lucientes, J. (2012). Host feeding patterns of *Culicoides* species (Diptera: Ceratopogonidae) within the Picos de Europa National Park in northern Spain. Bulletin of Entomological Research, 102(6), 692–697. 10.1017/S0007485312000284

9. Carpenter, S., Groschup, M. H., Garros, C., Felippe-Bauer, M. L., & Purse, B. V. (2013). Culicoides biting midges, arboviruses and public health in Europe. Antiviral Research, 100(1), 102–113. 10.1016/j.antiviral.2013.07.020

10. Carpenter, S., Wilson, A., & Mellor, P. S. (2009). *Culicoides* and the emergence of bluetongue virus in northern Europe. Trends in Microbiology, 17(4), 172–178. 10.1016/j.tim.2009.01.001

11. Casey-Bryars, M., Tratalos, J. A., Madden, J. M., & McGrath, G. (2025). Preparation for a potential outbreak of bluetongue virus in Ireland: Surveillance design to estimate local prevalence after an initial case detection. Irish Veterinary Journal, 78(1), 1–10. 10.1186/s13620-025-00315-1

12. Černý, O., Votýpka, J., & Svobodová, M. (2011). Spatial feeding preferences of ornithophilic mosquitoes, blackflies and biting midges. Medical and Veterinary Entomology, 25(1), 104–108. 10.1111/j.1365-2915.2010.00875.x

13. Chagas, C. R. F., Duc, M., Kazak, M., Valavičiūtė-Pocienė, K., Bukauskaitė, D., Hernández-Lara, C., & Bernotienė, R. (2024). High abundance of *Haemoproteus* parasites in *Culicoides* (Diptera: Ceratopogonidae), with confirmation of *Culicoides reconditus* as a new vector of these avian blood parasites. Insects, 15(3), 157. 10.3390/insects15030157

14. Collins, Á. B., Doherty, M. L., Barrett, D. J., & Mee, J. F. (2019). Schmallenberg virus: A systematic international literature review (2011-2019) from an Irish perspective. Irish Veterinary Journal, 72(1), 1–14. 10.1186/s13620-019-0147-3

15. Collins, Á. B., Mee, J. F., Doherty, M. L., Barrett, D. J., & England, M. E. (2018). Culicoides species composition and abundance on Irish cattle farms: implications for arboviral disease transmission. Parasites & vectors, 11(1), 472. 10.1186/s13071-018-3010-6

16. Curran, T., Browett, S., O’Neill, D., O’Hanlon, A., O’Reilly, C., Harrington, A., McDevitt, A., & O’Meara, D. (2022). One bat’s waste is another man’s treasure: a DNA metabarcoding approach for the assessment of biodiversity and ecosystem services in Ireland using bat faeces. Biodiversity and Conservation. 10.1007/s10531-022-02451-4

17. Dooley, J. J., Paine, K. E., Garrett, S. D., & Brown, H. M. (2004). Detection of meat species using TaqMan real-time PCR assays. Meat Science, 68(3), 431–438. 10.1016/j.meatsci.2004.04.010

18. Elbers, A. R. W., & Meiswinkel, R. (2015). *Culicoides* (Diptera: Ceratopogonidae) and livestock in the Netherlands: Comparing host preference and attack rates on a Shetland pony, a dairy cow, and a sheep. Journal of Vector Ecology, 40(2), 308–317. 10.1111/jvec.12169

19. England, M. E., Pearce-Kelly, P., Brugman, V. A., King, S., Gubbins, S., Sach, F., Sanders, C. J., Masters, N. J., Denison, E., & Carpenter, S. (2020). *Culicoides* species composition and molecular identification of host blood meals at two zoos in the UK. Parasites & Vectors, 13(1), Article 113. 10.1186/s13071-020-04018-0

20. Garros, C., Gardès, L., Allène, X., Rakotoarivony, I., Viennet, E., Rossi, S., & Balenghien, T. (2011). Adaptation of a species-specific multiplex PCR assay for the identification of blood meal source in *Culicoides* (Ceratopogonidae: Diptera): Applications on Palaearctic biting midge species, vectors of orbiviruses. Infection, Genetics and Evolution, 11(5), 1103–1110. 10.1016/j.meegid.2011.04.002

21. González, M. A., Goiri, F., Prosser, S. W. J., Cevidanes, A., Hernández-Triana, L. M., Barandika, J. F., Hebert, P. D. N., & García-Pérez, A. L. (2022). *Culicoides* species community composition and feeding preferences in two aquatic ecosystems in northern Spain. Parasites & Vectors, 15(1), Article 177. 10.1186/s13071-022-05297-5

22. Hadj-Henni, L., De Meulemeester, T., Depaquit, J., Noël, P., Germain, A., Helder, R., & Augot, D. (2015). Comparison of vertebrate cytochrome *b* and prepronociceptin for blood meal analyses in *Culicoides*. Frontiers in Veterinary Science, 2, Article 15. 10.3389/fvets.2015.00015

23. Isiye, E., Valcarcel Olmeda, A., Curran, T., O’Neill, D., de Waal, T., Barry, G., O’Hanlon, A., O’Shaughnessy, J., Keohane McCarthy, N., Vellinga, A., Jenkinson, A., Johnson, A., Barrett, D., Costello, S., Zintl, A., & O’Meara, D. (2025). Molecular characterisation of common *Culicoides* biting midges (Diptera: Ceratopogonidae) in Ireland. Parasites & Vectors, 18(1), Article 54. 10.1186/s13071-025-06754-7

24. Isiye, E., Olmeda, A. V., Curran, T., O’Neill, D., Waal, T. D., Barry, G., O’Hanlon, A., O’Shaughnessy, J., England, M., Zintl, A., & O’Meara, D. (2025). Design and optimisation of rapid real-time PCR assays for the detection of key *Culicoides* species. 10.1101/2025.09.26.678746

25. Kapustová, A., Kulich Fialová, M., Svobodová, M., & Brzoňová, J. (2025). Combining blood meal analysis and parasite detection yields a more comprehensive understanding of insect host feeding patterns. Parasites & Vectors, 18(1), Article 231. 10.1186/s13071-025-06931-8

26. Kasičová, Z., Schreiberová, A., Kimáková, A., & Kočišová, A. (2021). Blood meal analysis: Host-feeding patterns of biting midges (Diptera, Ceratopogonidae, Culicoides Latreille) in Slovakia. Parasite, 28, Article 58. 10.1051/parasite/2021058

27. Kazak, M., Valavičiūtė-Pocienė, K., Kondrotaitė, S., Duc, M., Bukauskaitė, D., Hernández-Lara, C., Bernotienė, R., & Chagas, C. R. F. (2024). *Culicoides* biting midges feeding behaviour as a key for understanding avian *Haemoproteus* transmission in Lithuania. Medical and Veterinary Entomology, 38(4), 530–541. 10.1111/mve.12752

28. Keven, J. B., Artzberger, G., Gillies, M. L., Mbewe, R. B., & Walker, E. D. (2020). Probe-based multiplex qPCR identifies blood-meal hosts in *Anopheles* mosquitoes from Papua New Guinea. Parasites & Vectors, 13(1), Article 55. 10.1186/s13071-020-3986-6

29. Kocher, T. D., Thomas, W. K., Meyer, A., Edwards, S. V., Pääbo, S., Villablanca, F. X., & Wilson, A. C. (1989). Dynamics of mitochondrial DNA evolution in animals: Amplification and sequencing with conserved primers. Proceedings of the National Academy of Sciences of the United States of America, 86(16), 6196–6200. 10.1073/pnas.86.16.6196

30. Lassen, S. B., Nielsen, S. A., & Kristensen, M. (2012). Identity and diversity of blood meal hosts of biting midges (Diptera: Ceratopogonidae: *Culicoides* Latreille) in Denmark. Parasites & Vectors, 5(1), Article 143. 10.1186/1756-3305-5-143

31. Martínez-de la Puente, J., Figuerola, J., & Soriguer, R. (2015). Fur or feather? Feeding preferences of species of *Culicoides* biting midges in Europe. Trends in Parasitology, 31(1), 16–22. 10.1016/j.pt.2014.11.002

32. Martínez-de la Puente, J., Martínez, J., Rivero-de Aguilar, J., Herrero, J., & Merino, S. (2011). On the specificity of avian blood parasites: Revealing specific and generalist relationships between haemosporidians and biting midges. Molecular Ecology, 20(15), 3275–3287. 10.1111/j.1365-294X.2011.05136.x

33. Martínez-de la Puente, J., Navarro, J., Ferraguti, M., Soriguer, R., & Figuerola, J. (2017). First molecular identification of the vertebrate hosts of *Culicoides imicola* in Europe and a review of its blood-feeding patterns worldwide: Implications for the transmission of bluetongue disease and African horse sickness. Medical and Veterinary Entomology, 31(4), 333–339. 10.1111/mve.12247

34. McGrath, G., More, S. J., & O’Neill, R. (2018). Hypothetical route of the introduction of Schmallenberg virus into Ireland using two complementary analyses. Veterinary Record, 182(8), 226. 10.1136/vr.104302

35. Meira, L., Costa, J., Villa, C., Ramos, F., Oliveira, M. B. P. P., & Mafra, I. (2017). EvaGreen real-time PCR to determine horse meat adulteration in processed foods. LWT, 75, 408–416. 10.1016/j.lwt.2016.08.061

36. Mignotte, A., Garros, C., Dellicour, S., Jacquot, M., Gilbert, M., Gardès, L., Balenghien, T., Duhayon, M., Rakotoarivony, I., de Wavrechin, M., & Huber, K. (2021). High dispersal capacity of *Culicoides obsoletus* (Diptera: Ceratopogonidae), vector of bluetongue and Schmallenberg viruses, revealed by landscape genetic analyses. Parasites & Vectors, 14(1), Article 22. 10.1186/s13071-020-04522-3

37. Mullins, J., Statham, M. J., Roche, T., Turner, P. D., & O’Reilly, C. (2010). Remotely plucked hair genotyping: A reliable and non-invasive method for censusing pine marten (Martes martes, L. 1758) populations. European Journal of Wildlife Research, 56(3), 443–453. 10.1007/s10344-009-0332-x

38. Ninio, C., Augot, D., Delecolle, J. C., Dufour, B., & Depaquit, J. (2011). Contribution to the knowledge of *Culicoides* (Diptera: Ceratopogonidae) host preferences in France. Parasitology Research, 108(3), 657–663. 10.1007/s00436-010-2110-9

39. Ninio, C., Augot, D., Dufour, B., & Depaquit, J. (2011). Emergence of *Culicoides obsoletus* from indoor and outdoor breeding sites. Veterinary Parasitology, 183(1-2), 125–129. 10.1016/j.vetpar.2011.07.020

40. Ogola, E. O., Bastos, A. D. S., Slothouwer, I., Getugi, C., Osalla, J., Omoga, D. C. A., Ondifu, D. O., Sang, R., Torto, B., Junglen, S., & Tchouassi, D. P. (2023). Viral diversity and blood-feeding patterns of Afrotropical *Culicoides* biting midges (Diptera: Ceratopogonidae). Frontiers in Microbiology, 14, 1325473. 10.3389/fmicb.2023.1325473

41. O’Meara, D. B., Sheehy, E., Turner, P. D., O’Mahony, D., Harrington, A. P., Denman, H., Lawton, C., MacPherson, J., & O’Reilly, C. (2014). Non-invasive multi-species monitoring: Real-time PCR detection of small mammal and squirrel prey DNA in pine marten (Martes martes) scats. Acta Theriologica, 59(1), 111–117. 10.1007/s13364-013-0155-8

42. Powell, C., Butler, F., & O’Reilly, C. (2019). The development of real-time PCR assays for species and sex identification of three sympatric deer species from noninvasive samples. Conservation Genetics Resources, 11(4), 465–471. 10.1007/s12686-018-1041-0

43. Purse, B. V., Carpenter, S., Venter, G. J., Bellis, G., & Mullens, B. A. (2015). Bionomics of temperate and tropical *Culicoides* midges: Knowledge gaps and consequences for transmission of *Culicoides*-borne viruses. Annual Review of Entomology, 60, 373–392. 10.1146/annurev-ento-010814-020614

44. R Core Team. (2023). R: A language and environment for statistical computing. R Foundation for Statistical Computing. https://www.r-project.org/

45. Reeves, L. E., Gillett-Kaufman, J. L., Kawahara, A. Y., & Kaufman, P. E. (2018). Barcoding blood meals: New vertebrate-specific primer sets for assigning taxonomic identities to host DNA from mosquito blood meals. PLoS Neglected Tropical Diseases, 12(8), e0006767. 10.1371/journal.pntd.0006767

46. Slama, D., Haouas, N., Mezhoud, H., Babba, H., & Chaker, E. (2015). Blood meal analysis of *Culicoides* (Diptera: Ceratopogonidae) in Central Tunisia. PLoS ONE, 10(3), e0120528. 10.1371/journal.pone.0120528

47. Statham, M. (2005). Development and use of DNA identification techniques for Irish mustelids (Doctoral thesis). Waterford Institute of Technology, Ireland.

48. Tarlinton, R., Daly, J., Dunham, S., & Kydd, J. (2012). The challenge of Schmallenberg virus emergence in Europe. The Veterinary Journal, 194(1), 10–18. 10.1016/j.tvjl.2012.08.017

49. Technelysium Pty Ltd. (2018). Chromas (Version 2.6.6) https://technelysium.com.au/wp/chromas/

50. Tomazatos, A., Jöst, H., Schulze, J., Spînu, M., Schmidt-Chanasit, J., Cadar, D., & Lühken, R. (2020). Blood-meal analysis of *Culicoides* (Diptera: Ceratopogonidae) reveals a broad host range and new species records for Romania. Parasites & Vectors, 13(1), Article 79. 10.1186/s13071-020-3938-1

51. Wernike, K., Fischer, L., Twietmeyer, S., & Beer, M. (2024). Extensive Schmallenberg virus circulation in Germany, 2023. Veterinary Research, 55(1), Article 134. 10.1186/s13567-024-01389-5

52. Wernike, K., Hoffmann, B., Link, E. K., Rotheneder, S., Eshak, M., Pfaff, F., Höper, D., & Beer, M. (2026). It happened again – Emergence of a Shamonda-like orthobunyavirus in Central Europe. bioRxiv. 10.64898/2026.08.06.743243

53. Wickham, H., Chang, W., Henry, L., Pedersen, T. L., Takahashi, K., Wilke, C., Woo, K., Yutani, H., & Dunnington, D. (2023). *ggplot2: Create elegant data visualisations using the grammar of graphics* [R package]. https://CRAN.R-project.org/package=ggplot2

